# *M. bovis* PPD Enhances Respiratory Bioenergetics of Human vs. Bovine Macrophages

**DOI:** 10.1101/2024.02.29.582730

**Authors:** Marie-Christine Bartens, Sam Willcocks, Dirk Werling, Amanda J. Gibson

**Affiliations:** Centre for Vaccinology and Regenerative Medicine, Department of Pathobiology and Population Science, Royal Veterinary College, UK; Department of Infection Biology, London School of Hygiene and Tropical Medicine, UK; Department of Life Sciences, Brunel University, UK; Department of Life Science, Aberystwyth University, UK

**Keywords:** Macrophage, Immunometabolism, Tuberculosis, Mycobacteria, BCG

## Abstract

The role of macrophage (MØ) cellular metabolism and reprogramming during TB infection is of great interest due to the influence of *Mycobacterium* spp. on MØ bioenergetics. Recent studies have shown that *M. tuberculosis* induces a TLR2-dependent shift towards aerobic glycolysis and metabolic reprogramming, comparable to the established LPS induced pro-inflammatory M1 MØ polarisation. Distinct differences in the metabolic profile of murine and human MØ indicates species-specific differences in bioenergetics. So far, studies examining the metabolic potential of cattle are lacking, thus the basic bioenergetics of bovine and human MØ were explored in response to a variety of innate immune stimuli. Cellular energy metabolism kinetics were measured concurrently for both species on a Seahorse XFe96 platform to generate bioenergetic profiles for the response to the bona-fide TLR2 and TLR4 ligands, FSL-1 and LPS respectively. Despite previous reports of species-specific differences in TLR signalling and cytokine production between human and bovine MØ, we observed similar respiratory profiles for both species. Basal respiration remained constant between stimulated MØ and controls, whereas addition of TLR ligands induced increased glycolysis. In contrast to MØ stimulation with *M. tuberculosis* PPD, another TLR2 ligand, *M. bovis* PPD treatment significantly enhanced basal respiration rates and glycolysis only in human MØ. Respiratory profiling further revealed significant elevation of ATP-linked OCR and maximal respiration suggesting a strong OXPHOS activation upon *M. bovis* PPD stimulation in human MØ. Our results provide an exploratory set of data elucidating the basic respiratory profile of bovine vs. human MØ that will not only lay the foundation for future studies to investigate host-tropism of the *M. tuberculosis* complex but may explain inflammatory differences observed for other zoonotic diseases.

**Highlights:**

- Similar baseline respiratory profiles for human and bovine macrophages
- *M. bovis* PPD treatment altered metabolic profile only in human MØ
- Strong OXPHOS activation upon *M. bovis* PPD stimulation only in human MØ

**Graphical Abstract:** Created with BioRender (www.biorender.com) by A. Gibson

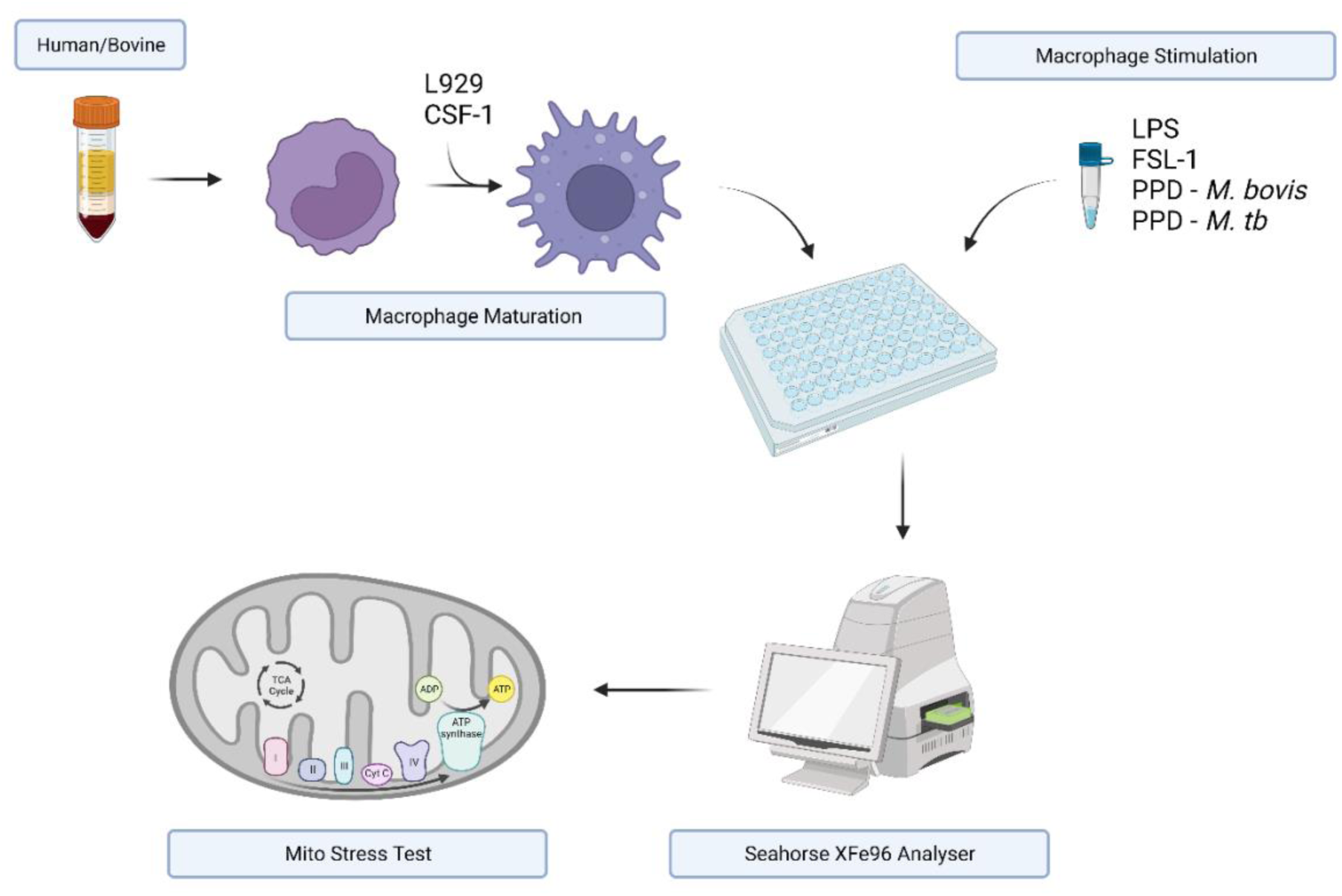

## Introduction

In the recent years, a growing interest in the cellular metabolism of innate immune cells has developed due to the understanding that changes in metabolic pathways of macrophages (MØ) in response to agonist stimulation impact on their phenotype and function (1–3). Numerous studies have emphasised that glycolysis, and therefore the provision of energy is crucial for immune cell function (2). Indeed, stimulation of MØ with various Pattern Recognition Receptors (PRR) ligands, most commonly Lipopolysaccharide (LPS), induces a metabolic shift from Oxidative Phosphorylation (OXPHOS) to glycolysis. This is considered a hallmark event in MØ activation, similar to the Warburg effect known in tumour cells (1, 4).

The Warburg effect occurs in tumour cells under normoxic conditions and glycolysis is the dominant metabolic pathway (4). During glycolysis, glucose is converted to pyruvate that enters the tricarboxylic acid cycle (TCA) cycle before being subsequently further metabolised by OXPHOS in the mitochondria (4). In tumour cells, pyruvate is metabolised to lactate instead of entering the TCA. Similar to this effect, activation of MØ induces a similar metabolic shift, with increased glycolysis, reduction in TCA cycle activity (5, 6) and increased lactate production and flux through the pentose phosphate pathway (PPP) (reviewed by Kelly and O’Neill (4)).

OXPHOS, like glycolysis, results in ATP production, though in significantly lesser in magnitude. However, glycolysis can be rapidly activated, which is important for MØ effector functions during pathogen infection, particularly host defence functions such as phagocytosis and production of inflammatory cytokines (2).

Indeed, altered MØ metabolism upon LPS plus Interferon-γ (IFN-γ) stimulation in comparison to interleukin-4 (IL-4) alone has formed the basis of priming MØ into either pro-inflammatory M1 MØ or anti-inflammatory M2 MØ (1, 2). This metabolic reprogramming leads to classically activated (M1) MØ being associated with host defence pathways, whereas alternately activated (M2) MØ promote T helper cell type 2 (TH2) driven immune responses and modulate repair processes (3). The main metabolic characteristics of M1 MØ are strongly enhanced glycolysis and impaired OXPHOS, similar to the Warburg effect described above (3). In combination with an enhanced PPP metabolism, this supports the resourcing of nucleotides for protein synthesis and increased Nicotinamide Adenine Dinucleotide Phosphate Hydrogen (NADPH) production for inflammatory MØ responses. Subsequent oxidation of NADPH results in the production (and release) of Reactive Oxygen Species (ROS), facilitating a direct bactericidal effect of MØ (7). To prevent hyper-inflammation of the tissue, NADPH is also used to generate glutathione and other antioxidants (2).

Furthermore, pyruvate generated by glycolysis fuels the TCA cycle and disrupts at the steps after citrate and succinate generation, leading to their subsequent accumulation, in pro-inflammatory M1 MØ (**Figure 1**) (4). The resulting citrate can be used for the synthesis of fatty acids, fatty acid derivates such as prostaglandins, the production of NO (8) and the generation of itaconic acid, a metabolite with direct anti-bactericidal effects against *M. tuberculosis* (MTB) (9). In addition, accumulated succinate stabilises hypoxia-inducible factor 1 α (HIF-1α), resulting in the maintenance of IL-1β production and thus supporting the generation of a pro-inflammatory response by MØ (8). HIF-1α is induced by hypoxia and inflammatory stimuli, and triggers and sustains glycolytic and pro-inflammatory pathways (1, 3). Indeed, HIF-1α activity is essential for the IFN-γ dependent MTB control. Lack of HIF-1α resulted in a strongly reduced pro-inflammatory cytokine and NO response and increased susceptibility to MTB infection in murine *in vitro* and *in vivo* models (9).

**Figure 1:**
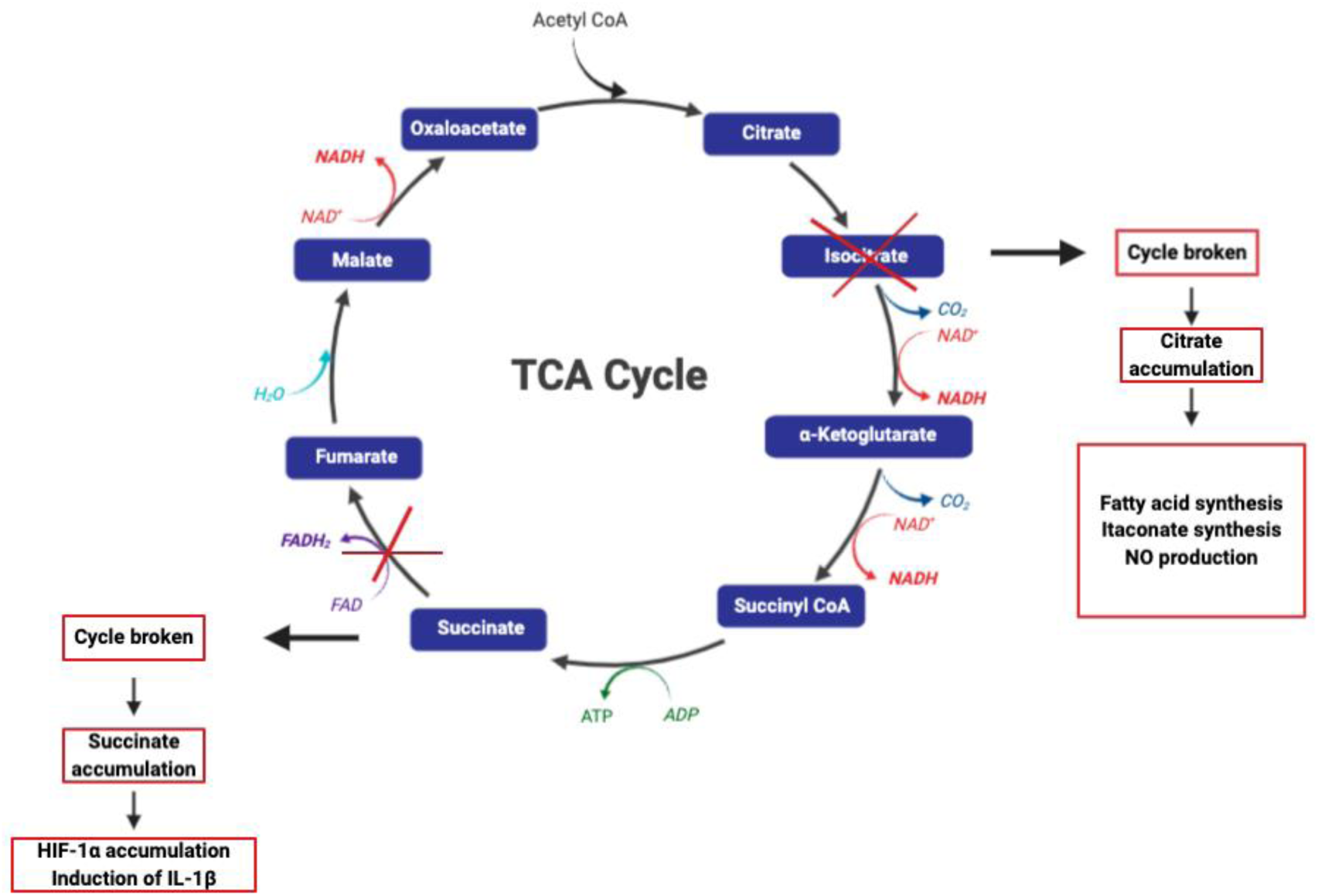
Disruption of the TCA cycle in M1 MØ. In M1-like MØ the TCA cycle is disrupted in two places — after citrate and after succinate leading to the accumulation of both metabolites. Image adapted from O’Neill *et al.* (3) and created with BioRender (www.biorender.com).

The vast majority of studies investigating immunometabolism have used LPS as a ligand (for example see ((10–16)). However, the role of MØ cell metabolism during tuberculosis (TB) infection has found great interest recently, recognising a major influence of MØ bioenergetics in the response to mycobacterial pathogens. Recent studies have shown that similar to LPS induced M1 MØ polarisation, MTB induces a metabolic shift in MØ towards aerobic glycolysis (9, 17–19). This shift has been shown to be TLR2 dependent (18) and furthermore HIF-1α coordinated in IFN-γ activated MØ (9), resulting in increased pro-inflammatory MØ effector function. However, this paradigm has been challenged lately as Cumming *et al.* (20) reported a downregulation of both, OXPHOS and glycolysis, upon live MTB infection in human MØ, suggesting the induction of a quiescent energy phenotype by live MTB in primary cells. Shi *et al.* (19) also recently reported a biphasic dynamic of MØ metabolism with an early phase, characterised by M1 MØ polarisation, but a late adaptation post 24 h with transition from glycolysis to OXPHOS, indicating a subsequent downregulation of MØ pro-inflammatory and anti-bactericidal responses. Thus, MØ immunometabolism is clearly an emerging field and further studies elucidating the complexity of metabolic changes induced by pathogens in the host are indicated to improve our understanding.

Most investigations into cellular immunometabolism have been conducted in the murine model or by using cell lines. Very recently, distinct differences in the metabolic profile of murine and human MØ have been identified (16), suggesting species-specific differences. So far, studies examining the metabolic potential of bovine MØ are lacking, thus the current study investigates the basic respiratory parameters and bioenergetics of bovine and human MØ were comparatively explored in response to a variety of ligands.

## Materials and Methods

### Cell Culture

#### Isolation of bovine peripheral blood mononuclear cells (PBMCs)

Blood for peripheral blood mononuclear cells (PBMC) isolation and subsequent MØ generation was collected by puncture of the jugular vein from clinically healthy pure-breed pedigree Holstein Friesian (HF) and Brown Swiss (BS) cows housed at the RVC Bolton Park Farm (Hertfordshire, UK) and Cancourt Farm (Wiltshire, UK). All procedures were carried out under the Home Office license (PPL7009059) approved by the RVC’s Ethics and Welfare Committee. For biological assays, blood was drawn into sterile glass vacuum bottles containing 10% acid citrate dextrose (ACD) as anticoagulant and isolated as previously described (21, 22). Serum was collected using vacutainers from the same animals.

#### Maturation and culture of bovine *ex-vivo* derived macrophages

PBMCs were isolated as previously described (21, 22) To derive MØ, thawed PBMCs were set up in 10 x 10 cm dishes in MØ cell culture media, supplemented with 10% L929 fibroblast cell line supernatant as source of M-CSF (produced by the Werling group, RVC) at 1 x 10^6^ cells ml^-1^ in a total volume of 20 ml and incubated at 37°C with 5% CO2. Media was replaced after three days and cells were harvested after 6 days. Cells were scraped of the dishes using cell scrapers (Greiner, UK) and cold PBS. After assessing cell viability by Trypan Blue (Sigma, UK) exclusion, cells were seeded at 1 x 10^6^ ml^-1^ in 96-well plates for further assays.

#### Isolation, maturation and culture of human PBMCs

Human blood was collected from healthy donors in 50 ml EDTA tubes at LSHTM under a CREB with ethics approval (No 2019 1916-3). Sex and *M. bovis* BCG vaccination status of the donors was recorded. PBMCs were isolated in the same manner as described above for the bovine PBMCs. To derive MØ from frozen PBMC stocks, cells were treated as described for the bovine MØ.

#### Extracellular flux assay

To accurately investigate the bioenergetic function of bovine and human MØ with an extracellular flux analyser (Seahorse Bioscience, Inc, USA), both cell types were characterized according to the manufacturer’s recommended basal and test conditions (23).

Initial experiments were conducted with an 8-well Seahorse XFp extracellular flux analyser (Seahorse Bioscience, Inc, USA) to determine cell seeding density and FCCP concentration (Supplementary Data). All following experiments investigating metabolic parameters of both cell types upon ligand stimulation, were conducted with a 96-well Seahorse XFe extracellular flux analyser (Seahorse Bioscience, Inc, USA).

#### Investigation of metabolic parameters

Key metabolic parameters of human and bovine MØ were determined in real time by measuring oxygen consumption rate (OCR) and Extracellular Acidification Rate (ECAR) using a Seahorse XFe 96-well extracellular flux analyzer (Agilent, USA). Briefly, *ex-vivo* derived bovine and human MØ were matured as described above and seeded at a density of 1.5 x 10^5^ cells in 180 µl in XFe cell culture microplates (Agilent, USA). Cells were stimulated with LPS (1 ng ml^-1^; Invivogen, USA), FSL-1 (100 ng ml^-1^; Invivogen, USA), recombinant bovine (rbo) TGFβ1 (10 ng ml^-1^, National Institute of Biological Standards and Control (NIBSC), UK) at, PPD *M. bovis* (1 µg ml^-1^, NIBSC, UK) or *M. bovis* BCG Pasteur at an MOI of 10 for 24 h incubation. Following the incubation, cells were washed, and MØ cell culture media was replaced with FCS-and bicarbonate-free DMEM medium supplemented with 4.5 mg ml^-1^ D-glucose and 2 mM glutamine (Agilent, USA) for another 60 min incubation at 37°C without CO2. The XFe96 sensor cartridge was hydrated overnight prior to the assay and used to calibrate the analyser. Compounds of the Mito Stress Test kit (Seahorse Bioscience, Inc, USA) target components of the electron transport chain (ETC) were prepared according to the manufacturers’ instructions. After calibration, the cell culture plate was loaded and basal OCR and ECAR were recorded following by sequential addition of the compounds of the Cell Mito Stress Test kit (23). Firstly, oligomycin (inhibitor of ATP synthase) was added to reach a final concentration of 1 μM, followed by FCCP (uncoupling agent) at a final concentration of 2.0 μM and subsequently rotenone/antimycin A (inhibitors of complex I and complex III of the respiratory chain, respectively) to reach a final concentration of 0.5 μM per wells (See Supplementary Figure 1). Data was recorded in wave software and exported to Excel and GraphPad Prism (Dotmatics, V8.4.3). Parameters are then extrapolated using the multi-report generator files from Agilent Technologies, which automatically calculate proton leak and spare respiratory capacity using measured parameters during the assay, such as ATP production, maximal respiration, and non-mitochondrial respiration.

##### 1.5.2 Statistical Analysis

Statistical analysis and graphs were generated using GraphPad Prism software package (Version 7, GraphPad Inc., USA). All results were checked for normal distribution and equal variance assumption and are presented as mean +/- standard deviation. Datasets that passed normality tests and equal variance assumptions, but appeared to have outliers, were log transformed and re-analysed to verify no change in statistical results. Output of statistical analysis using log transformed datasets is listed in the Appendix. Datasets that were additionally log transformed to account for outliers are clearly marked within the manuscript. Statistical significance was defined as p<0.05(*), p <0.01(**), p<0.001(***) and p < 0.0001(****).

## Results

### 2-Deoxy-D-glucose potently inhibits glycolysis in bovine and human MØ

Using 2-Deoxy-D-glucose (2-DG) to inhibit the first step of glycolysis has been shown to be a useful tool to examine glycolytic parameters more closely (4, 6, 16). Here, we first examined the ability of 2-DG to control MØ metabolism and to examine whether its effects were similar in both species. We observed that 2-DG decreased OCR (Figure 2A and C) and strongly decreased glycolysis in MØ of both species to a greater extent than control ligands (LPS and FSL-1), which were used for comparison in the same experiments (Figure 2B and D).

**Figure 2:**
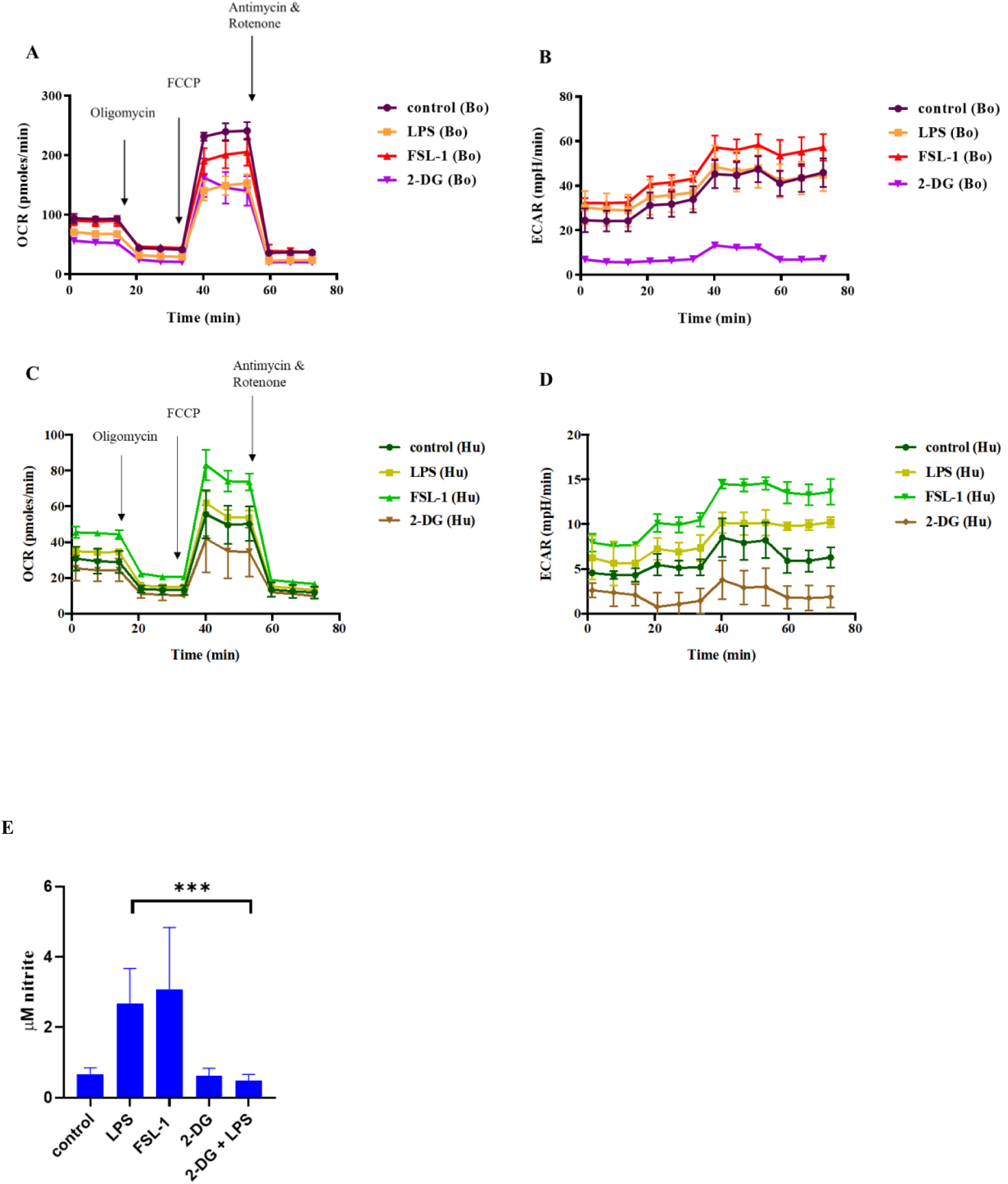
Respiratory profiles of bovine and human MØ to 2-DG stimulation. Representative plot of OCR **(A and C)** and ECAR **(B and D)** of a bovine MØ (Bo) (n=1, **A and B)** and a human (Hu) MØ sample (n=1, **C and D**) upon stimulation with 1 mM 2-DG (Sigma-Aldrich, UK) for 24 h subjected to a Cell Mito Stress test (Agilent Technologies, USA) and measured with the Seahorse XFe extracellular flux analyser (Agilent Technologies, USA) over 75 min. Following measurement of basal respiration, OCR and ECAR are recorded after injection of 1 μM Oligomycin, followed by 2.0 μM FCCP injection and 0.5 μM of antimycin A and rotenone injection (injection time points are indicated by arrows). MØ generated from cattle (n=4) were stimulated with either 1 mg/ml LPS (Invivogen, USA), 100 ng/ml FSL-1 (Invivogen, USA), 1 mM 2-DG (Sigma-Aldrich, UK) or 1 mg/ml LPS+ 1 mM 2-DG for 24 h. Thereafter, supernatants were collected before cells were subjected to Mito Stress test, and frozen until measurement of NO production by Griess assay (Promega, UK) **(E)**. Graphs and statistical analysis (one-way ANOVA) were prepared in GraphPad Prism V8 (GraphPad Inc., USA). All samples were run in duplicates and mean +/- SD are shown.

A shift in MØ metabolism towards increased glycolysis was reported to be important for MØ effector function, since inhibition of glycolysis by 2-DG has been shown to decrease the inflammatory response (reviewed by Kelly and O’Neill (4)). Indeed, NO production was significantly impaired upon addition of 2-DG to LPS stimulation on cattle MØ (Figure 2E) measured using supernatants from stimulated MØ just before performing a Cell Mito Stress test. Furthermore, a statistically significant difference between the cattle breeds upon FSL-1 stimulation was observed.

### Bovine and human MØ exhibit similar mitochondrial bioenergetics pattern upon LPS stimulation

Most studies have assessed MØ cell metabolism in M1/M2 MØ, where the M1 phenotype was generated upon LPS stimulation. Thus, we examined cellular bioenergetics in bovine and human MØ in response to the TLR4 agonist LPS.

Levels of OCR are an indicator of OXPHOS. Initially basal respiration was measured before the injection of cell Mito stress test compounds to determine specific respiratory parameters. Basal respiration was similar in LPS-stimulated MØ and their corresponding controls for both species (Figure 3A). ECAR, an indicator of glycolysis, was mildly elevated in both species upon LPS stimulation. Upon LPS stimulation, the overall respiratory profile differed between control and stimulated groups for both species, though only very mildly for basal respiration, resulting in a minor decrease of OCR for BS and human MØ, whereas glycolysis was increased for all groups (Figure 3B). An example of a respiratory profile for a bovine and human sample is shown in Figure 4. While the magnitude of difference varied greatly between the two species, the overall trend remained the same for each species.

**Figure 3:**
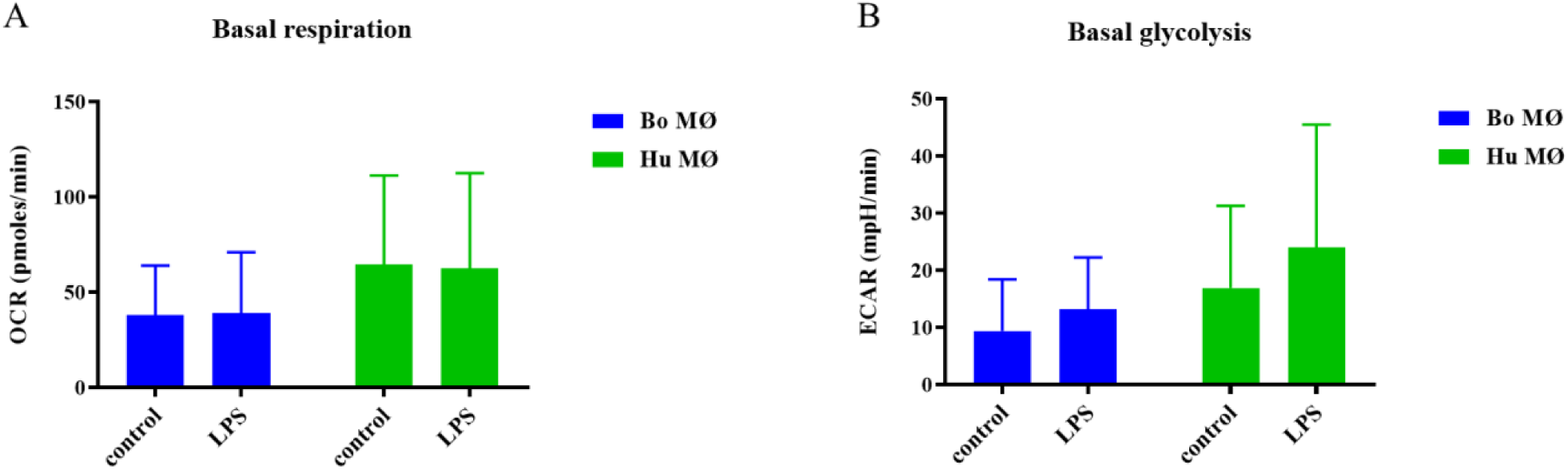
Basal respiration and glycolysis of bovine and human MØ to LPS stimulation. Basal respiration and glycolysis of bovine (Bo) (n=8) and human (Hu) MØ (n=4) upon stimulation with 1μg/ml LPS (Invivogen, USA) for 24 h measured using the Seahorse XFe extracellular flux analyser (Agilent Technologies, USA) prior to a Cell Mito stress test (Agilent Technologies, USA). Values are the mean of three independent experiments with three technical repeats each and are shown as +/- SD. Statistical analysis was performed using paired Student’s t-test relative to controls in GraphPad Prism V8 (GraphPad Inc., USA). Statistically significant difference is indicated by asterisk (* =p<0.05).

**Figure 4:**
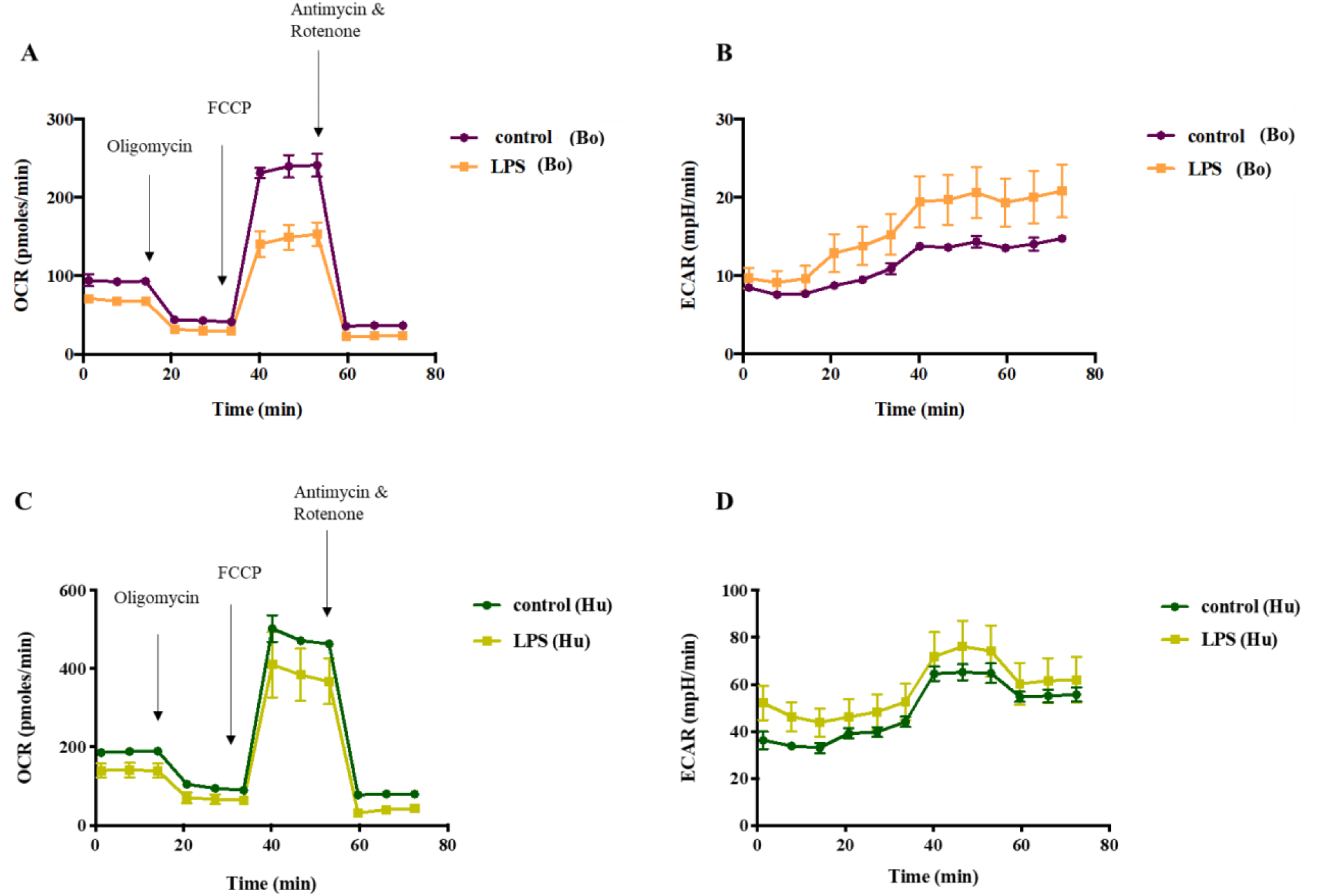
Respiratory profiles of bovine and human MØ to LPS stimulation. A representative plot of OCR **(A and B)** and ECAR **(C and D)** of a bovine (Bo) and human (Hu) MØ sample upon stimulation with 1μg/ml LPS (Invivogen, USA) for 24 h subjected to a Cell Mito stress test (Agilent Technologies, USA) and measured using a Seahorse XFe extracellular flux analyser (Agilent Technologies, USA) over 75 min is shown. Following measurement of basal respiration, OCR and ECAR are recorded after injection of 1 μM Oligomycin, followed by 2.0 μM FCCP injection and 0.5 μM of antimycin A and rotenone (injection time points are indicated by arrows). Graphs are displayed in GraphPad Prism V8 (GraphPad Inc., USA).

Examining the corresponding respiratory parameters (Figure 5), LPS stimulation did not alter the response greatly for human MØ, though a tendency towards a minimal increase of non-mitochondrial respiration (Figure 5D) and decline of proton leak was observed (Figure 5E). Cattle responses to LPS showed minimally impaired respiratory parameters upon LPS stimulation, (Figure 5B-D), although this reduction was not statistically significant. The strong individual variation between samples was reflected in all respiratory parameters.

**Figure 5:**
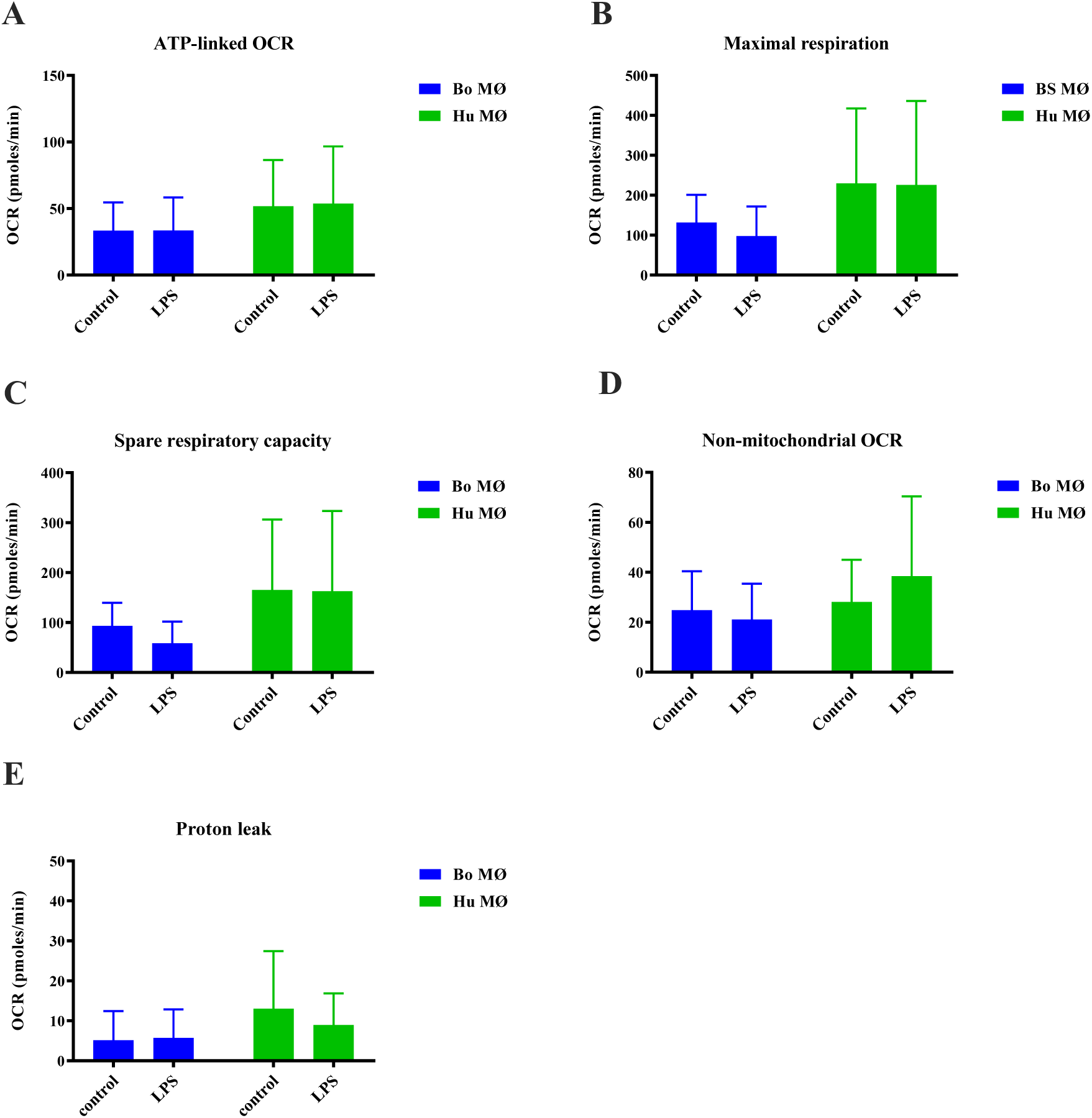
Respiratory parameters of bovine and human MØ to LPS stimulation. Respiratory parameters of bovine (Bo, n=8) and human (Hu) MØ (n=4) upon stimulation with 1 μg/ml LPS (Invivogen, USA) for 24 h subjected to a Cell Mito stress test (Agilent Technologies, USA) measured with the Seahorse XFe extracellular flux analyser (Agilent Technologies, USA). Values are the mean of three independent experiments with three technical repeats each and are shown as +/- SD. Statistical analysis was performed using paired Student’s t-test relative to controls in GraphPad Prism V8 (GraphPad Inc., USA).

In summary, considering the data obtained from all animals, it appeared that mitochondrial bioenergetics was not significantly altered in either species upon LPS stimulation. However, it is noteworthy that basal parameters were consistently the highest for human MØ. However, no further analysis of this difference was performed, as the aim of the study was to examine cellular bioenergetics of MØ in response to ligand stimulation.

### Bovine and human MØ exhibit similar mitochondrial bioenergetics upon FSL-1 stimulation

Having observed a similar mitochondrial bioenergetic profile upon LPS stimulation between the two species, we next investigated the response to FSL-1, a TLR2 agonist. Breed specific responses to FSL-1 has been described by us and others (24, 25). A similar profile of mitochondrial bioenergetics as observed upon LPS stimulation was detected. Basal respiration equalled among controls and stimulated groups for both species and glycolysis rates were elevated in the stimulated groups (Figure 6). This increase was significantly increased for BS MØ (Figure 6B). Interestingly, respiratory parameters were similar between control and stimulated groups for MØ generated from humans and cattle upon FSL-1 stimulation, in contrast to their decreased response upon LPS stimulation (Figure 7A-E).

**Figure 6:**
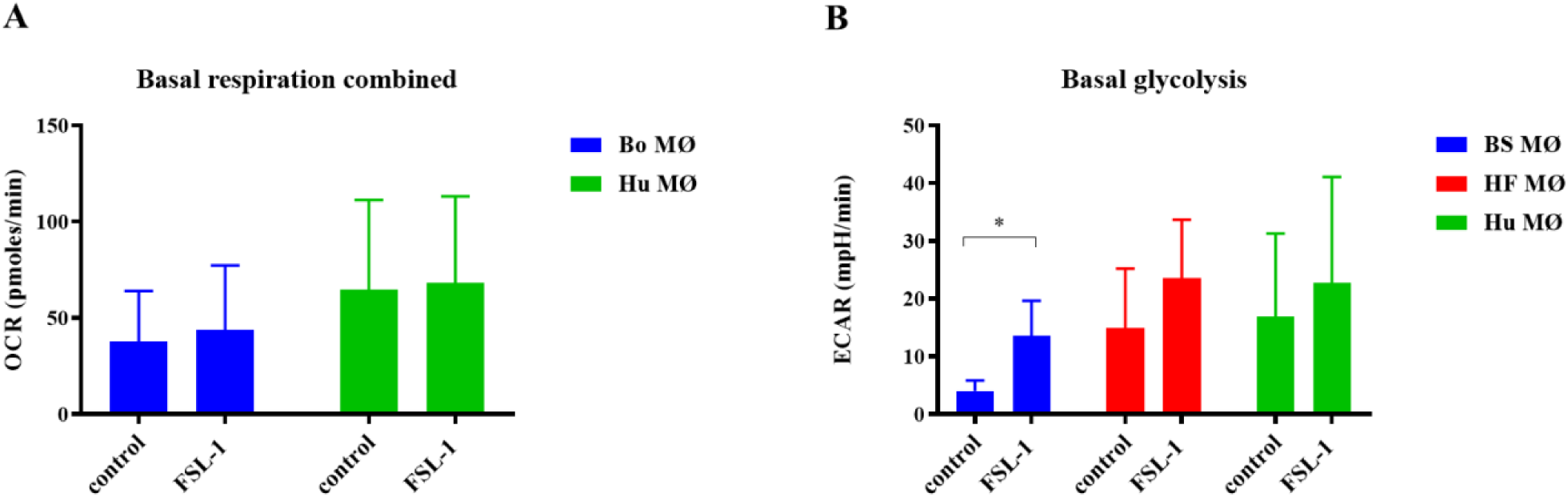
Basal respiration and glycolysis of bovine and human MØ to FSL-1 stimulation. Basal respiration and glycolysis of bovine ((Bo, n=4 per breed) and human (Hu) MØ (n=4) upon stimulation with 100 ng/ml FSL-1 (Invivogen, USA) for 24 h measured with the Seahorse XFe extracellular flux analyser (Agilent Technologies, USA) prior to a Cell Mito stress test (Agilent Technologies, USA). Values are the mean of three independent experiments with three technical repeats each and are shown as +/- SD. Statistical analysis was performed using paired Student’s t-test relative to controls in GraphPad Prism V8 (GraphPad Inc., USA). Statistically significant difference is indicated by asterisk (* =p<0.05).

**Figure 7:**
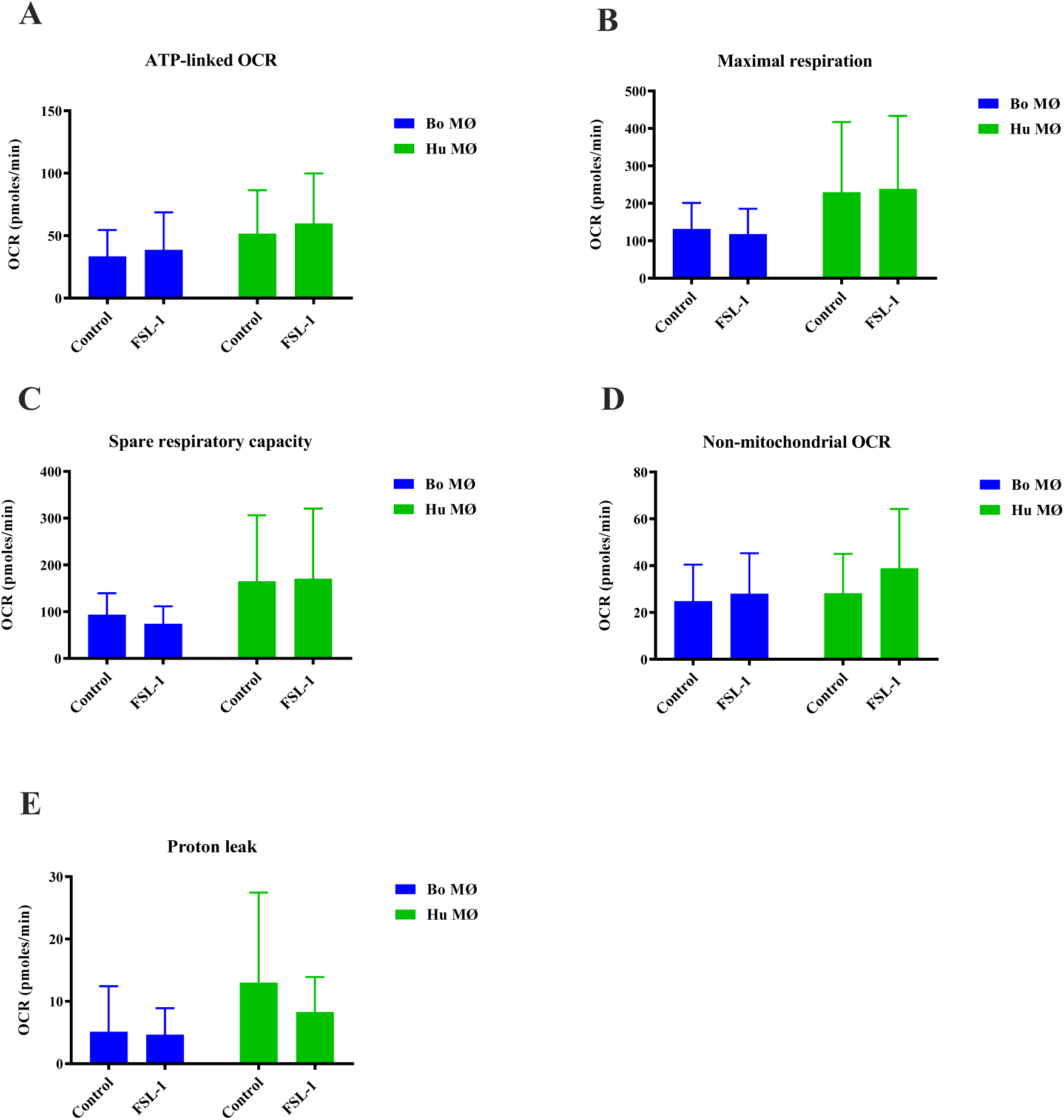
Respiratory parameters of bovine and human MØ to FSL-1 stimulation. Respiratory parameters of bovine (Bo, n=8) and human (Hu) MØ (n=4) upon stimulation with 100 ng/ml FSL-1 (Invivogen, USA) for 24 h measured with the Seahorse XFe extracellular flux analyser (Agilent Technologies, USA) prior to a Cell Mito stress test (Agilent Technologies, USA). Values are the mean of three independent experiments with three technical repeats each and are shown as +/- SD. Statistical analysis was performed using paired Student’s t-test relative to controls in GraphPad Prism V8 (GraphPad Inc., USA).

### Enhanced activation of respiratory parameters upon PPD of *M. bovis* stimulation in human MØ

As experiments with the extracellular flux analyser were only possible under Biosafety level 2 laboratory conditions, purified protein derivative (PPD) derived from *M. bovis* (NIBSC, UK) was used as a substitute rather than using fully virulent *M.bovis*. Interestingly, basal respiration was minimally increased in bovine MØ, but strongly and significantly increased in human MØ (Figure 8A). Similarly, basal glycolysis was not significantly elevated for bovine MØ, but for human MØ only (Figure 8B), indicating a species difference upon stimulation with PPD of *M. bovis*. Furthermore, with the exemption of proton leak, all respiratory parameters were elevated in human MØ which was found to be significant for ATP-linked OCR and maximal respiration (Figure 9A-E), suggesting a strong OXPHOS activation upon PPD of *M. bovis* stimulation in human MØ (Figure 9A-E).

**Figure 8:**
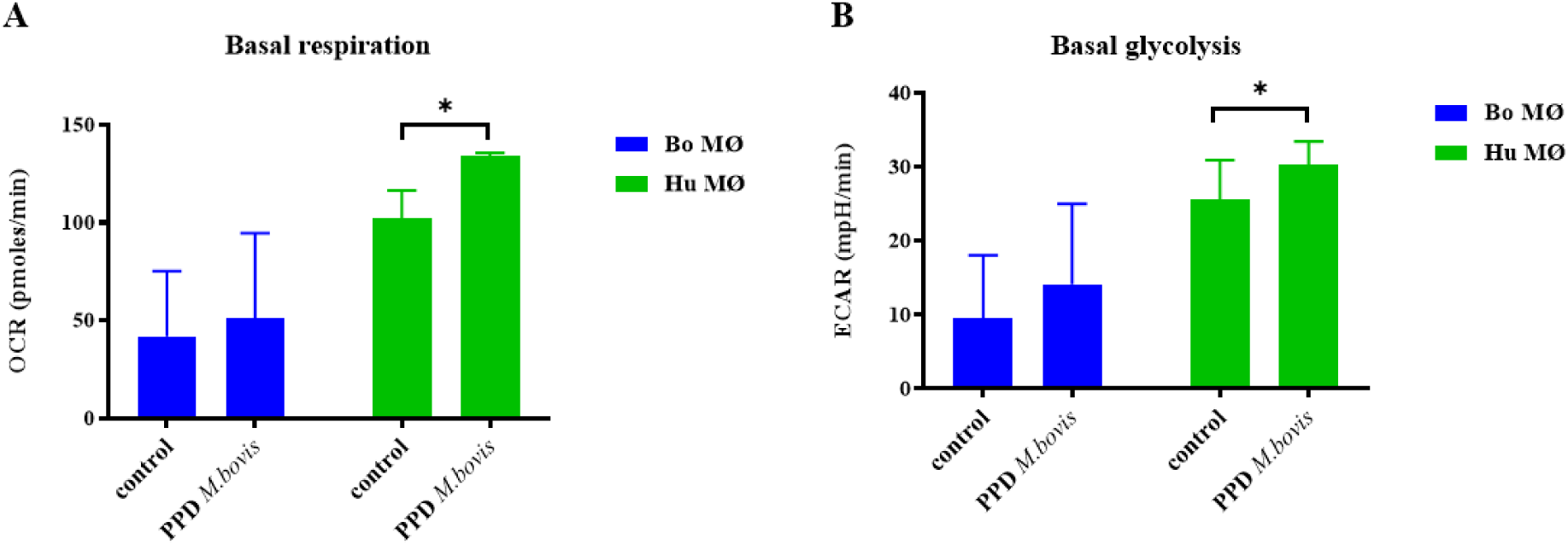
Basal respiration and glycolysis of bovine and human MØ to PPD of *M. bovis* stimulation. Basal respiration and glycolysis of bovine (Bo, n=8) and human (Hu) MØ (n=4) upon stimulation with 1 μg/ml PPD of *M. bovis* (NIBSC, UK) for 24 h measured with the Seahorse XFe extracellular flux analyser (Agilent Technologies, USA) prior to a Cell Mito Stress test (Agilent Technologies, USA). Values are of the mean of three independent experiments with three technical repeats each and are shown as +/- SD. Statistical analysis was performed using paired Student’s t-test relative to controls in GraphPad Prism V8 (GraphPad Inc., USA). Statistically significant differences are indicated by asterisk (*=p<0.05).

**Figure 9:**
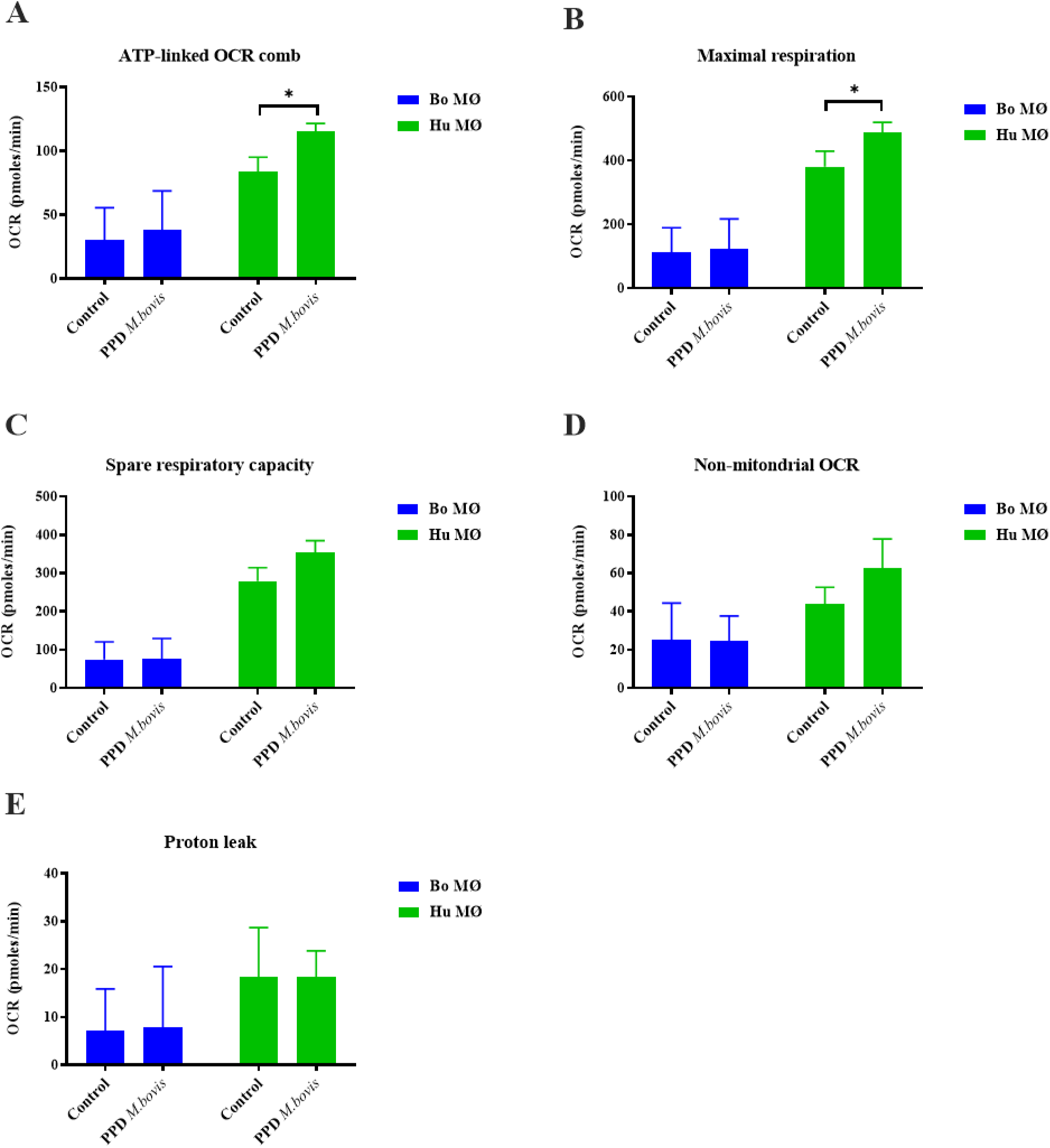
Respiratory parameters of bovine and human MØ to PPD of *M. bovis* stimulation. Respiratory parameters of bovine (Bo, n=8) and human (Hu) MØ (n=4) upon stimulation with 1μg/ml PPD of *M. bovis* (NIBSC, UK) for 24 h measured with the Seahorse XFe extracellular flux analyser (Agilent Technologies, USA) prior to a Cell Mito Stress test (Agilent Technologies, USA). Values are the mean of three independent experiments with three technical repeats each and are shown as +/- SD. Statistical analysis was performed using paired Student’s t-test relative to controls in GraphPad Prism V8 (GraphPad Inc., USA). Statistically significant differences are indicated by asterisk (*=p<0.05).

### Bovine and human MØ exhibit similar mitochondrial bioenergetics upon PPD of M. *tuberculosis* stimulation

To assess mitochondrial bioenergetics in response to *M. tuberculosis* (MTB), PPD derived from MTB (NIBSC, UK) was also used as comparable substitute for live virulent MTB. As observed before for PPD of *M. bovis*, human MØ showed a tendency for a higher basal respiration and glycolysis upon PPD of MTB stimulation, however compared to PPD derived from *M. bovis* this was found not to be significant (Figure 10). For MØ generated from cattle, measurements in controls equalled those in stimulated cells for both parameters (Figure 10). Respiratory parameters for bovine and human MØ showed similar levels in control and stimulated groups, besides an increase in non-mitochondrial respiration in human MØ (Figure 11A-E).

**Figure 10:**
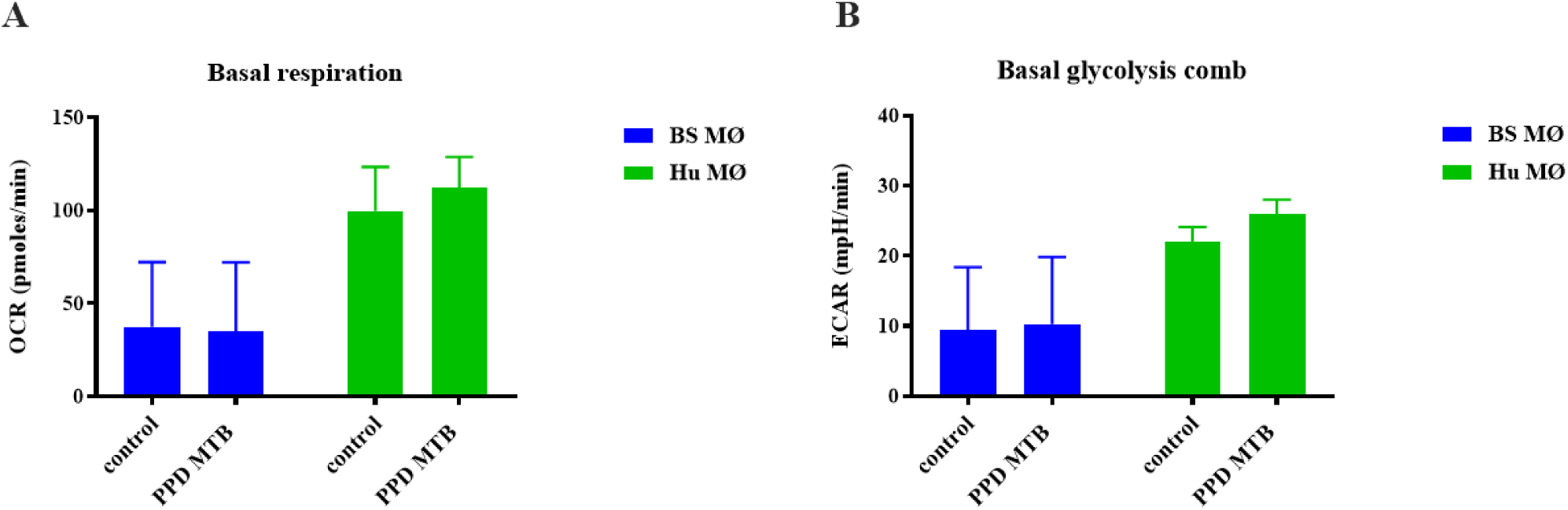
Basal respiration and glycolysis of bovine and human MØ to PPD of MTB stimulation. Basal respiration and glycolysis of bovine (Bo, n=6) and human (Hu) MØ (n=3) upon stimulation with 1 μg/ml PPD of MTB (NIBSC, UK) for 24 h measured with the Seahorse XFe extracellular flux analyser (Agilent Technologies, USA) prior to a Cell Mito Stress test (Agilent Technologies, USA). Values are the mean of three independent experiments with three technical repeats and are shown as +/- SD. Statistical analysis was performed using paired Student’s t-test relative to controls in GraphPad Prism V8 (GraphPad Inc., USA).

**Figure 11:**
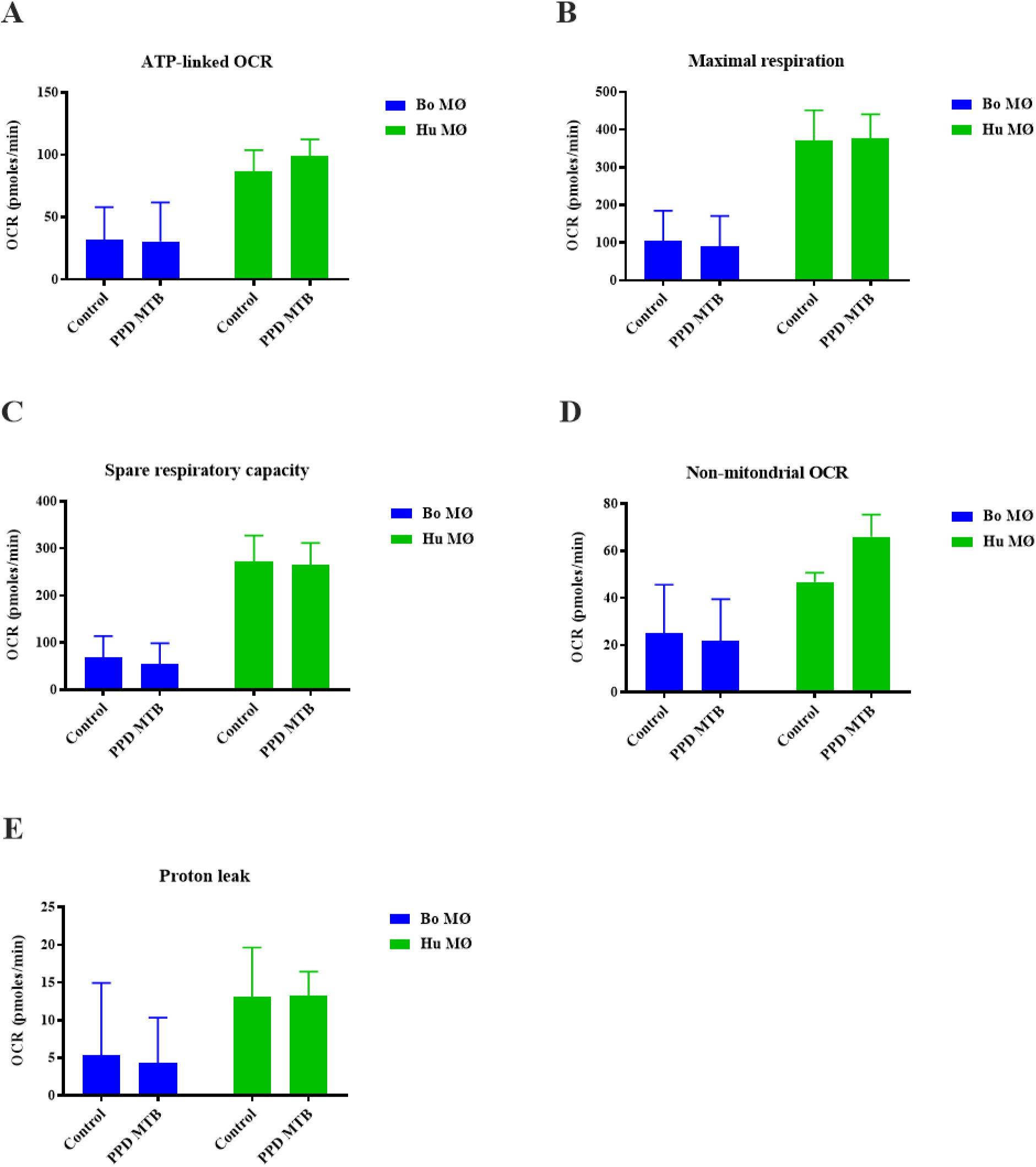
Respiratory parameters of bovine and human MØ to PPD of MTB stimulation. Respiratory parameters of bovine (Bo, n= 4) and human (Hu) MØ (n=2) upon stimulation with 1 μg/mL PPD of MTB (NIBSC, UK) for 24 h subjected to a Cell Mito Stress test (Agilent Technologies, USA) measured with the Seahorse XFe analyser (Agilent Technologies, USA). Values are the mean of three independent experiments with three technical repeats each and are shown as +/- SD. Statistical analysis was performed using paired Student’s t-test relative to controls in GraphPad Prism V8 (GraphPad Inc., USA).

## Discussion

The aim of the experiments described in the present study was to explore some basic immune-metabolic functions of bovine and human MØ in response to a variety of ligands. Most studies in this field have been conducted using murine MØ or cell lines to investigate metabolic changes, however functional differences between MØ types and species have been reported (ref needed). No prior investigations into the cellular bioenergetics of bovine MØ have been made so far. However, given the in general reduced response to TLR ligands seen in bovine MØ(26–28), we wanted to assess whether a similar phenomenon could be observed comparing the metabolism of bovine MØ and human MØ using a Seahorse extracellular flux analyser and a Cell Mito Stress test (both Agilent Technologies, USA) which allowed the determination of key parameters of mitochondrial respiration in both species.

Overall, there was a trend for increased glycolysis in response to all ligands (with exception of the glycolysis inhibitor 2-DG). An increase in glycolysis upon stimulation with LPS, but also to mycobacteria is considered a hallmark event in activated MØ (1, 4). This increase in glycolytic pathways allows for rapid energy production and triggers host defence pathways such as pro-inflammatory cytokine and effector molecule expression (2–4) For instance, metabolic reprogramming has been shown to be essential in control of mycobacterial infection (9, 17–19). Inhibition of glycolysis by 2-DG, which was used in a control experiment here, has been shown to enhance mycobacterial growth, suggesting that glycolysis is required for limiting MTB growth (29). Here NO production, an essential antimicrobial, was also found to be ameliorated upon 2-DG treatment in all cell types analysed indicating that depleting glycolysis would have similar effects for mycobacterial control in human and bovine MQ.

When comparing respiratory parameters between the bovine and human species, no distinct differences were detected. This differed from results published in a recent study, reporting significant differences in the metabolic profile of human and murine MØ upon LPS stimulation, showing impaired mitochondrial bioenergetics in the murine samples (16). In contrast, we observed similar mitochondrial bioenergetics in MØ generated from both species in response to LPS, FSL-1, PPD of MTB and 2-DG. Interestingly, several authors have described different responses of either human and bovine MØ exposed to either mycobacterial species (25) as well differences how bacterial infection is dealt with in cattle (30, 31). These previous observations triggered our experiments to assess whether observed changes might be at least in part explained by metabolic changes in MØ from either species. Like our data, a stronger activation of mitochondrial respiration was observed to PPD of *M. bovis* in human MØ. Cummings *et al.* (20) also observed an increase in the overall OCR measured in the Cell Mito Stress test and some respiratory parameters such as maximal respiration and spare respiratory capacity upon *M. bovis* BCG stimulation of human MØ in contrast to exposure to live and dead MTB. It could be hypothesised that *M. bovis* activates human MØ more strongly, which is further supported by the absence of a significantly stronger activation upon PPD of MTB stimulation in human MØ. In their study (20), MTB drastically decreased MØ respiratory parameters which was found to be MOI dependent. However, it has to be kept in mind that Cummings *et al.* (20) used live bacteria, in contrast to the present study using PPD derivative only. Indeed, the authors (20) observed distinct differences in the respiratory capacity of MØ to live and dead MTB and *M. bovis* BCG, and additionally these were dependent on the MØ type (cell line vs primary cells) and dose of infection.

The overall response pattern did not vary significantly between breeds or species. Highest basal values were measured in human MØ and lowest for MØ generated from BS cattle. Higher basal values of human MØ were also found in the study by Vijayan *et al.* (16) in comparison to murine bone marrow-derived MØ. However, this was likely just a reflection of the MØ type used under specific cell culture conditions, whereas the focus of the present study was to examine the response of the bovine and human MØ generated in the same manner to identical ligands.

In general, the source of MØ generation, their method of culture and maturation, differential stimulation periods and dose of infections have been shown to strongly impact the metabolic profile of MØ and subsequently the outcome of the response. In the present study a consistent approach was used, as MØ of both species were generated in the same manner and always stimulated for 24 h. Nonetheless, through this approach some time-dependent metabolic changes as reported by Shi *et al*. (19) may have been missed. Additionally, the MØ lineage of the same species seemed to impact on their metabolic profile. Huang *et al.* (29) found differences in the metabolism of interstitial and alveolar MØ during MTB infection, with the former showing a higher glycolytic activity and the latter skewed to fatty acid oxidation.

The results presented here were exploratory, elucidating a basic respiratory profile of bovine MØ in comparison to human MØ. Further investigations into specific metabolic pathways using live bacteria are indicated to allow for further assumptions. However, considering these preliminary results, it can be hypothesised that bovine MØ have similar bioenergetic profile to human MØ upon stimulation with the most commonly used PAMP, such as LPS and others used herein, and thus might serve as a useful comparative tool to study MØ metabolism further.

### Similar MØ bioenergetics between both species

There is growing evidence linking MØ metabolism to the production of inflammatory mediators (6). Upon stimulation with various ligands, MØ undergo metabolic reprogramming, resulting in impaired OXPHOS and increased glycolysis (2, 4) and the latter has been found essential for pro-inflammatory MØ function (4). Overall, only a mild inhibition of mitochondrial respiration in response to agonists was found in this study, though an increase of ECAR, the indicator for glycolysis, was more pronounced in both species to all agonists. Interestingly, inhibition of OXPHOS was found not essential for MØ to sustain inflammatory polarization, contrary to glycolysis whose upregulation is needed for inflammatory function and cell survival (32).

Recently, NO has been demonstrated to be a central modulator of this metabolic switch. NO can act as inhibitor of Complex I and reversibly, Complex IV of the ETC (15, 32). Increased levels of NO as observed for mycobacterial infection as well as in supernatants from cells subsequently used for metabolism assays (Figure 2E), may explain the decreased mitochondrial respiration upon agonist stimulation observed in some samples.

Through the upregulation of glycolysis, TCA cycle metabolites citrate and succinate are accumulated. Succinate accumulation leads to the stabilisation of HIF-1α and subsequent IL-1β and ROS production. Elevated succinate levels are attributed to itaconate, a metabolite with direct bactericidal effect against MTB (33) and that has recently been shown to be modulated by NO (32). Furthermore, citrate accumulation has been shown to induce NO production (2), further supporting the observed elevated NO levels.

## Abbreviations

2-DG: 2-Deoxy-D-Glucose
ACD: Acid Citrate Dextrose
ANOVA: Analysis of Variance
ATP: Adenosine Tri-phosphate
BCG: *M. bovis* strain Bacillus-Calmette Guérin
BS: Brown Swiss
DMEM: Dulbecco’s Minimal Essential Medium
ECAR: Extracellular Acidification Rate
EDTA: Ethylenediaminetetraacetic Acid
ETC: Electron Transport Chain
FCCP: Carbonyl Cyanide-p-(trifluoromethoxy) phenylhydrazone
FSL-1: TLR-2/6 ligand representing N-terminus of LP44 from *Mycoplasma salivarium*
HF: Holstein Friesian
HIF-1α: Hypoxia Inducible Factor-1α
HSD: Honestly Significant Difference (for Tukey’s HSD test)
IFN-γ: Interferon-γ
IL-4: Interleukin-4
LPS: Lipopolysaccharide
LSHTM: London School of Hygiene and Tropical Medicine
M1: Type 1 Macrophages
M2: Type 2 Macrophages
MM: Macrophage Media
MØ: Macrophage
MOI: Multiplicity of Infection
MTB: M. tuberculosis
mTOR: Mammalian Target of Rapamycin
NADPH: Reduced Nicotinamide Adenine Dinucleotide Phosphate
NIBSC: National Institute of Biological Standards and Control
NO: Nitric Oxide
OCR: Oxygen Consumption Rate
OXPHOS: Oxidative Phosphorylation
PBMC: Peripheral Blood Mononuclear Cells
PBS: Phosphate Buffered Saline
PPD: Purified Protein Derivative
PPP: Pentose Phosphate Pathway
PRR: Pattern Recognition Pathway
ROS: Reactive Oxygen Species
TB: Tuberculosis
TCA: Tricarboxylic Acid Cycle
TGF-β1: Transforming Growth Factor - β1
TH1: T helper 1 cells
TH2: T helper 2 cells
TLR: Toll-like Receptor

