## Supplementary Material for "*M. bovis* PPD Enhances Respiratory Bioenergetics of Human vs. Bovine Macrophages"

**Supplementary Information**

### Introduction

#### Extracellular flux assay

Mitochondria are the central energy hub of immune cells as they generate ATP through OXPHOS (26). Here, MØ energy metabolism in bovine and human MØ was investigated. To determine the effect of various stimuli on MØ OXPHOS and to measure corresponding key respiratory parameters, the extracellular flux analyser including a Cell Mito Stress Test kit (both Agilent Technologies, USA) was used. This allowed for the real-time measurement of oxygen consumption rate (OCR) and extracellular acidification rate (ECAR) in live cells upon sequentially injecting compounds to determine specific parameters. A schematic illustration of this process is shown in Figure S1. The key respiratory parameters analysed were the basal respiration and glycolysis that are measured as initial OCR and ECAR before injection of Cell Mito Stress compounds. Furthermore, ATP-linked OCR which was measured upon injection of oligomycin, a complex V inhibitor of the electron transport chain (ETC), is an indicator of respiration needed for ATP synthesis. Maximal respiration was calculated upon injection of (trifluoromethoxy) phenylhydrazone (FCCP), an uncoupling agent that disrupts the mitochondrial membrane potential and allows for maximum oxygen consumption. Additionally, spare respiratory capacity was calculated as the difference between maximal and basal respiration, a parameter that assesses the cell’s capacity to respond to increased energy demand in conditions of stress. Upon injection of the last compound of the Mito Stress test, antimycin A and rotenone, which inhibit the remaining complexes of the ETC, non-mitochondrial respiration was obtained, due to the general shutdown of OXPHOS. Finally, proton leak was calculated as the difference between ATP-linked OCR and non-mitochondrial respiration, that can be either a regulatory mechanism of ATP production or a sign of mitochondrial damage.

### Materials and Methods

#### Determination of the optimal cell seeding density

In a first step, the optimal seeding concentration for bovine and human macrophages was determined as the provided cell culture miniplate for all used extracellular flux analyser (XFp and XFe) differs from a standard 96 well plate (approximately 40% of the surface area). Thus, bovine and human macrophages were seeded at different concentrations, ranging from 1.0-1.5x $10^5$ cells per well, and visually assessed for confluency after seeding, a one-hour incubation time as well as after some washing steps. An optimal seeding density was determined at 1.5x$10^5$ cells per well for both cell types, resulting in 90% confluency. Thus, this seeding density was used for all subsequent experiments.

**Determination of FCCP concentration**

Upon manufacturer recommendation, a 5-point titration curve was performed to identify the optimal carbonyl cyanide 4-(trifluoromethoxy) phenylhydrazone (FCCP) concentration which results in the maximal oxygen consumption rate (OCR) for both cell types, bovine and human macrophages. For this, the cell culture plate was divided into two groups: A low concentration range (0.125 µM, 0.25 µM, 0.50 µM) and a high concentration range (0.50 µM, 1.0 µM, 2.0 µM). Each group was treated with 1 µM oligomycin first, followed by three serial injections of FCCP at the above-mentioned concentrations. Oxygen consumption of the cells' response to the five different doses of FCCP was recorded for 100 min and FCCP concentration that induced the highest OCR (2.0 µM for both cell types) was subsequently chosen for all future experiments.

### Results

**Optimisation of extracellular flux assay conditions**

To accurately investigate the bioenergetic function of bovine and human MØ with the extracellular flux analyser, MØ of both species were characterized using the manufacturer's recommended basal and test conditions. For these initial experiments, the Seahorse extracellular flux analyser XFp (allows measurement in 8-well plate format) (Agilent Technologies, USA) was used whereas for all subsequent experiments (from 7.3.5 onwards) the Seahorse extracellular flux analyser XFe (allows measurement in 96-well plate format) (Agilent Technologies, USA) was used.

**Determination of the optimal cell seeding density**

In a first step, the optimal seeding concentration for bovine and human MØ was determined. The provided cell culture miniplate differs from a standard 96 well plate as it has approximately

40% of the surface area. Thus, bovine and human MØ were seeded at different concentrations ($1.0 - 2.0 \times 10^5$ cells per well) and visually assessed for confluency after seeding and washing steps over 24 h. For MØ of both species, an optimal seeding density was determined at $1.5 \times 10^5$ cells per well (data not shown). This resulted in 90% confluency which generated desirable measurement ranges with both extracellular flux analysers and thus was used in all subsequent experiments.

**Determination of the optimal FCCP titration**

To identify the optimal FCCP concentration which results in the maximal OCR, a 5-point titration with serial FCCP injections was performed upon manufacturer's recommendation. Cells were divided in low and high FCCP concentration groups and OCR was recorded for at least 80 min. For MØ of both species, maximal OCR was determined upon 2.0 μM FCCP stimulation and therefore used in all subsequent experiments (Figure S2, A and B).

**Figures**

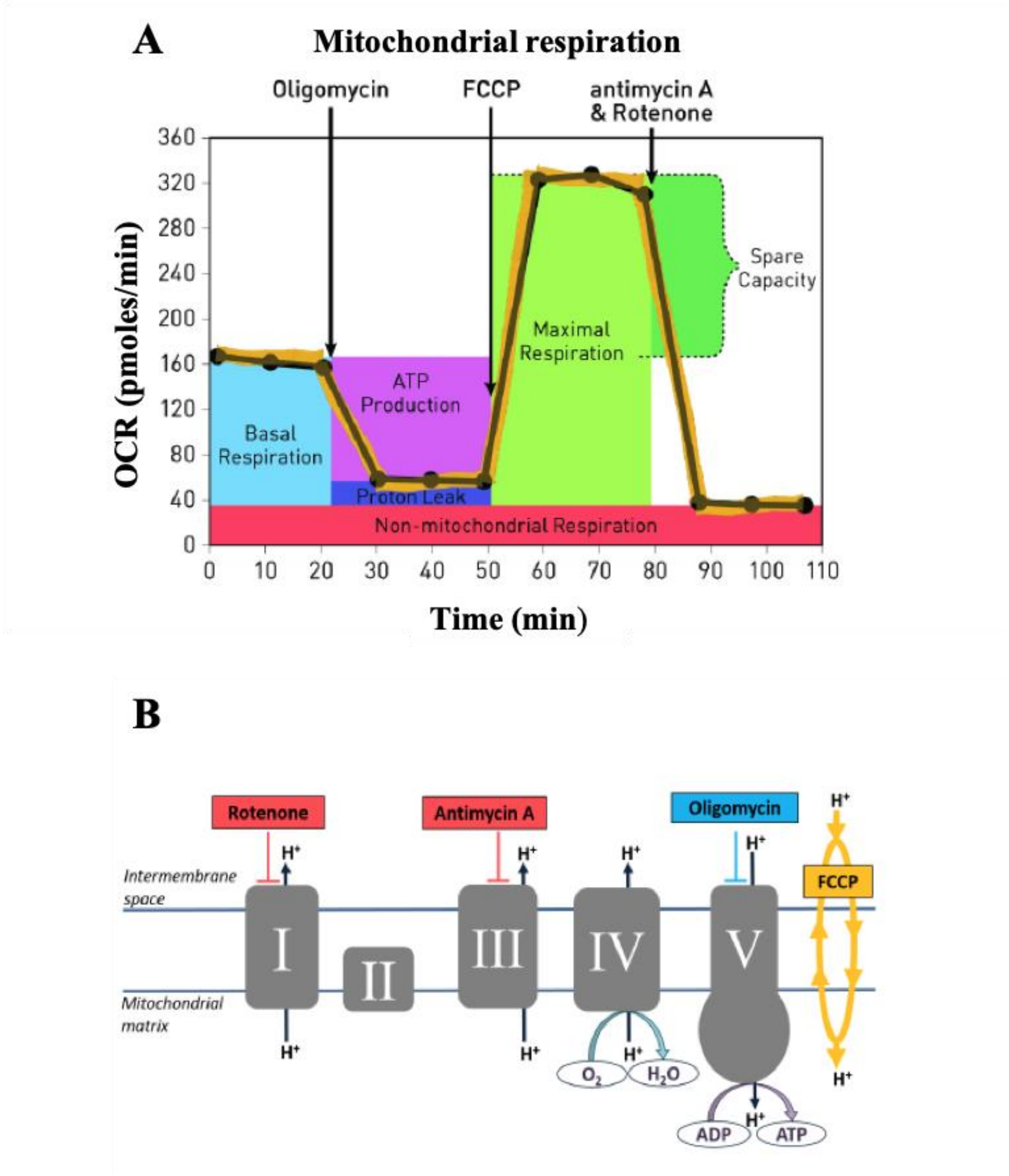


**Figure S11**: **Representative illustration of Cell Mito Stress test**

Illustration of the workflow including the sequential compound injection of oligomycin, FCCP and antimycin A and rotenone during the Cell Mito Stress test when measured by a Seahorse extracellular flux analyser (Agilent Technologies, USA) (A). The specific complexes of the ETC targeted by the compounds are displayed in (B) (27).

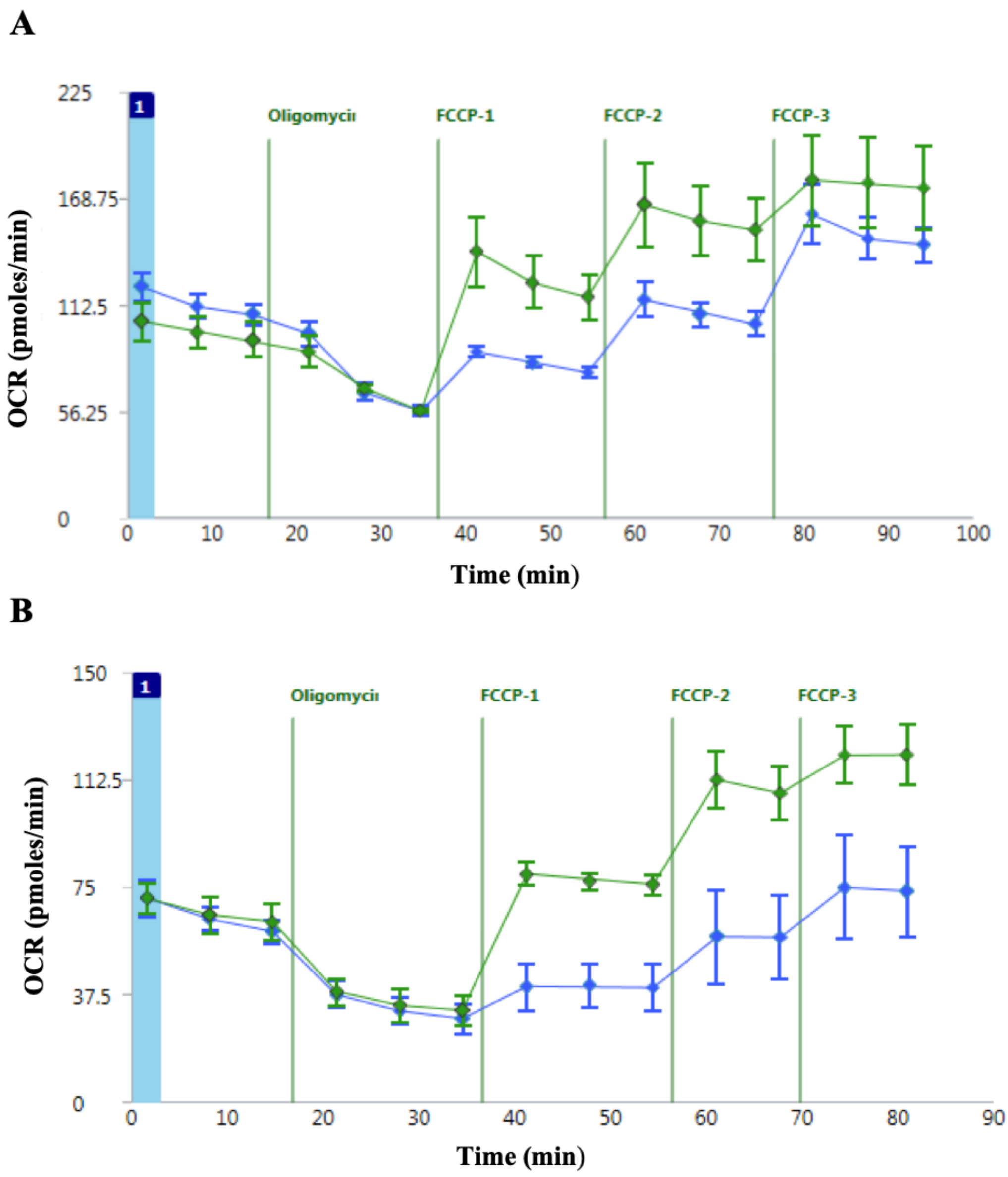


**Figure S2: Titration curves to determine optimal FCCP concentration**

Two 5-point titration curves were performed to identify the FCCP concentration which results in the maximal oxygen consumption rate (OCR) of human (A) and bovine (B) MØ. For this, the MØ cell culture plate was divided into two groups: A low concentration range (FCCP-1: 0.125 µM, FCCP-2: 0.25 µM, FCCP-3: 0.50 µM) shown in blue and a high concentration range (FCCP-1: 0.50 µM, FCCP-2: 1.0 µM, FCCP-3: 2.0 µM) shown in green. Each group was treated with 1 µM oligomycin first, followed by three serial injections of FCCP at the above-mentioned concentrations and at time points indicated on the graphs. OCR of the MØ response to the five different doses of FCCP was recorded for at least 80 min using a Seahorse XFp extracellular flux analyser (Agilent Technologies, USA). Samples were run in triplicates and are shown as mean +/- SD.
